# SpectralTDF: transition densities of diffusion processes with time-varying selection parameters, mutation rates, and effective population sizes

**DOI:** 10.1101/029736

**Authors:** Matthias Steinrücken, Ethan M. Jewett, Yun S. Song

## Abstract

In the Wright-Fisher diffusion, the transition density function (TDF) describes the time-evolution of the population-wide frequency of an allele. This function has several practical applications in population genetics, and computing it for biologically realistic scenarios with selection and demography is an important problem. We develop an efficient method for finding a spectral representation of the TDF for a general model where the effective population size, selection coefficients, and mutation parameters vary over time in a piecewise constant manner. The method, called spectralTDF, is available at https://sourceforge.net/projects/spectraltdf/.

## 1. Introduction

The transition density function (TDF) of the Wright-Fisher diffusion describes the time-evolution of the frequency of an allele (Ewens 2004). The TDF is useful for understanding the effects of demography, mutation, and selection on genetic variation, and it is a key component of a number of methods for inferring selection coefficients (Williamson *et al*. 2004, Bollback *et al*. 2008, Steinrücken *et al*. 2014), predicting allele fixation times (Waxman 2011), and computing population genetic statistics such as the site frequency spectrum (Živković *et al*. 2015).

Most existing approaches for computing the TDF assume either restrictive models of dominance (Kimura 1955; 1957) or selective neutrality (Shimakura 1977, Griffiths 1979, Vogl 2014a), or are computationally slow for selection strengths commonly observed in biological data (Barbour *et al*. 2000). However, Song and Steinrücken (2012), Steinrücken *et al*. (2013) recently developed a numerically stable and computationally efficient method for finding a spectral representation of the TDF for a general selection model in the case of constant parameters (population size, mutation rates, and selection coefficients). Despite the utility of this new approach, assuming that model parameters remain constant over time is often too restrictive for biological applications (Siepielski *et al*. 2009).

Živković *et al*. (2015) have extended the spectral method of Song and Steinrücken (2012) to handle piecewise-constant population size functions. However, their approach requires a restricted model of selection in which the fitness of a homozygote is twice that of a heterozygote (i.e., additive or genic selection). Furthermore, selection parameters are assumed to remain constant over time and the model does not allow for recurrent mutations.

Here, we present the first method for computing the TDF under arbitrary models of dominance and recurrent mutation while allowing selection parameters, mutation rates, and effective population sizes to change over time in a piecewise constant manner.

## 2. Model and approach

We consider a biallelic locus with two alleles, *A*_0_ and *A*_1_, evolving in a single panmictic population. In the corresponding Wright-Fisher diffusion, *X_t_* denotes the frequency of allele *A*_1_ at time *t*, measured continuously in units of generations. We assume that either *X*_0_ is given or the distribution of *X*_0_ is specified. The effective population size, mutation rates, and selection parameters are assumed to be constant within each of *K* disjoint epochs. As illustrated in Figure 1, the *k*th epoch has effective size *N_k_* (diploid individuals) and duration *τ_k_*. Epoch boundaries are denoted by *t*_0_; *t*_1_, …, *t_K_*, with 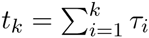.

**Figure 1.**
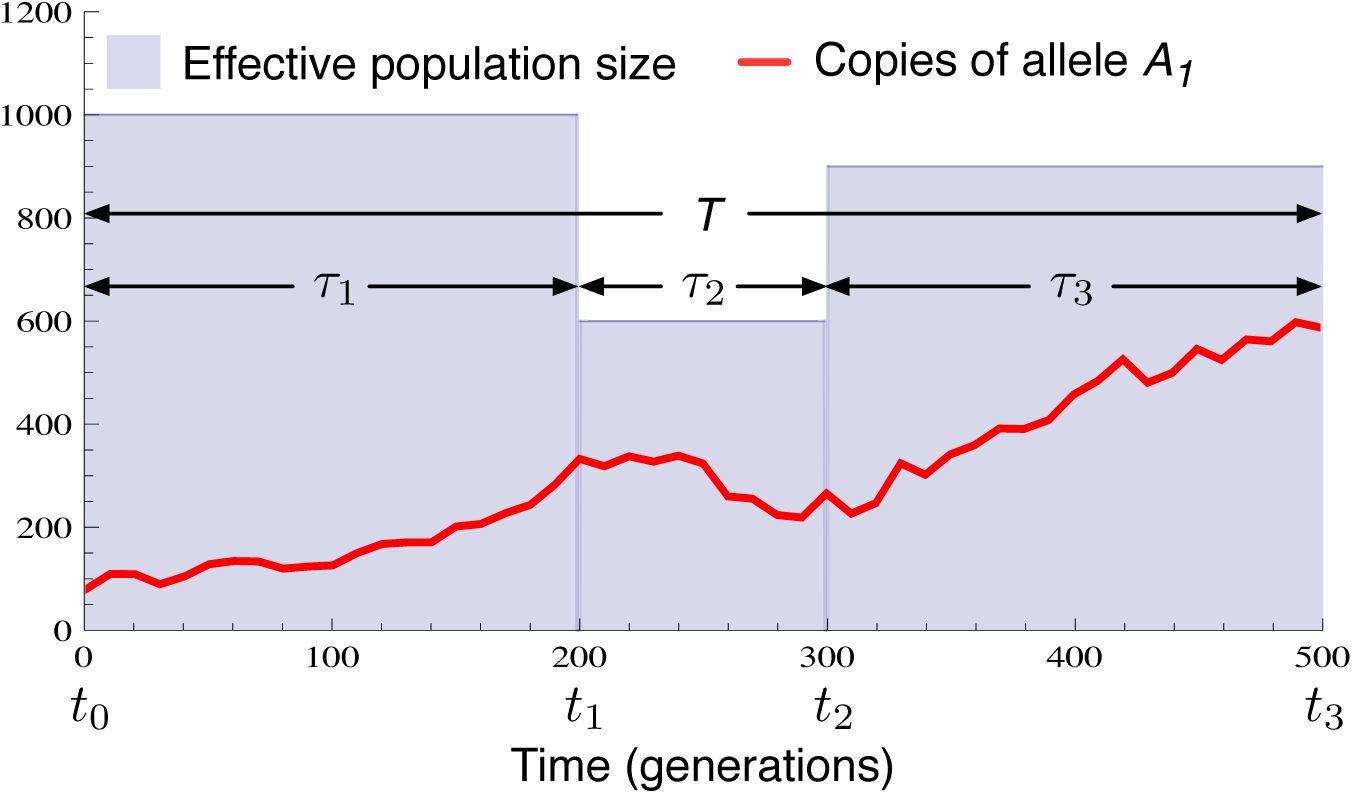
Diagram of the model. A population has constant size in each of *K* epochs (*N*_1_ = 1000, *N*_2_ = 600, *N*_3_ = 900). An allele, *A*_1_, at a locus of interest evolves over time, subject to pressures of mutation and selection that are constant within each epoch.

Within the *k*th epoch, the per-generation probability that a copy of allele *A*_0_ mutates to allele *A*_1_ is *a_k_*, and the per-generation probability that a copy of allele *A*_1_ mutates to allele *A*_0_ is *b_k_*. In addition, selection acts in such a way that the relative fitness of an individual carrying *i* copies of allele *A*_1_ is 1 + *s_ki_* (*i* = 1, 2).

Within each epoch, *k*, a spectral representation of the TDF, *p_k_*(*t*; *x*, *y*), can be obtained by employing the framework of Song and Steinrücken (2012). The challenge in computing the TDF for the full model with *K* epochs lies in knitting together the expressions for the densities *p_k_*(*t*; *x*, *y*) across the different epochs. We first review the derivations of Song and Steinrücken (2012) and Steinrücken *et al*. (2014) of the TDF in a single epoch of constant size. We then discuss our efficient polynomial interpolation method for knitting together the TDF across epochs of different constant sizes.

### 2.1. The TDF and its generalization in a single epoch of constant size.

We wish to compute the TDF in the *k*th epoch, where the density is defined by 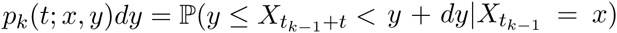, for *t* ∈ [*t_k_*_–1_, *t_k_*). We are also interested in the generalization, 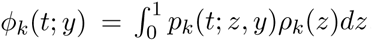, which extends the TDF to the case of a general initial density *ρ_k_* rather than a point mass at *x*.

In an epoch of constant size *N_k_*, let *α_k_* = 4*N_k_a_k_*, *β_k_* = 4*N_k_b_k_*, *σ_k_*,_1_ = *N_k_s_k_*, _1_, and *σ_k_*_,2_ = *N_k_s_k_*_,2_ denote population-size scaled versions of the per-generation mutation parameters (*a_k_, b_k_*) and selection coefficients (*s_k_*_,1_, *s_k_*_,2_). The Kolmogorov backward operator 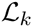 for the epoch is the second-order linear differential operator given by

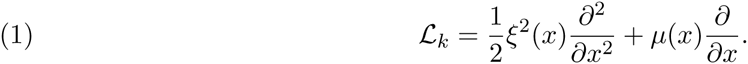

In Equation (1), the quantity

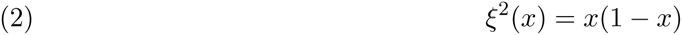

captures the contribution from genetic drift and the term

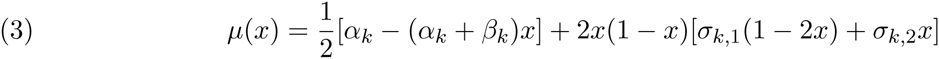

captures the contribution from recurrent mutation and selection. The TDF is the solution of the Kolmogorov backward equation

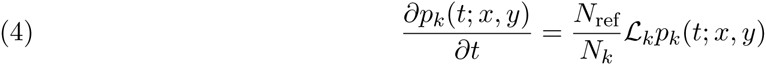

satisfying specified boundary conditions [see Song and Steinrücken (2012), p. 119, for a discussion of the boundary conditions]. We measure time in units of 2*N*_ref_ generations in all epochs, where *N*_ref_ is the size of a fixed reference population. For ease of interpretation, we choose *N*_ref_ = 1/2 so that time is measured in units of generations in all epochs.

Song and Steinrücken (2012) derived a formula for the TDF by obtaining a solution of the backward Equation (4) in the form of the infinite series

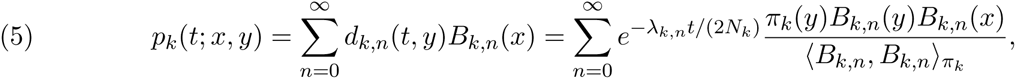

where 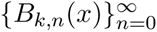 is the set of eigenfunctions of 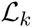 with associated eigenvalues 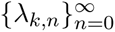 (Section 2.2.1) and the function *π_k_*(*y*) is given by

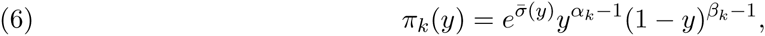

where 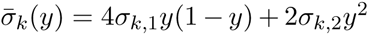. The inner product 〈*f*, *g*〉*_ω_* with respect to a weight function *ω*(*x*) in Equation (5) is defined for two functions *f* and *g* on an interval [*a*, *b*] by

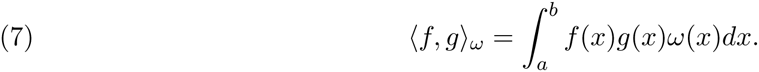

In Equation (5), the inner product 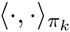 is taken over the interval [0; 1] with respect to *π_k_*(*y*). Equation (5) can be thought of as a function in either the initial frequency, *x*, or the final frequency, *y*.

### 2.2. The generalized TDF.

The generalization *ϕ_k_*(*t*; *y*) = 〈*p_k_*(t; ·, *y*), *ρ_k_*〉 of Equation (5) to the case in which the initial probability density is given by *ρ_k_* (*x*) is easily obtained by noting that, viewed as an expansion in the basis functions 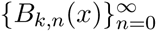, the partial sum 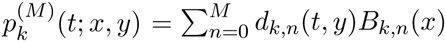 converges strongly to *p_k_*(*t*; *x*, *y*), from which it follows that

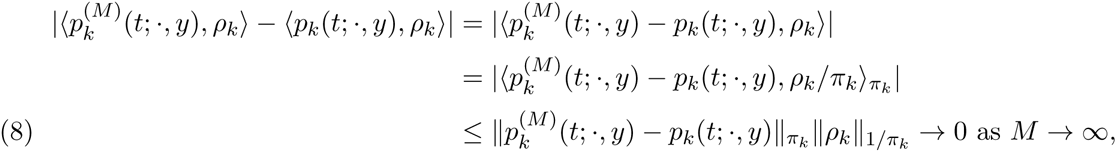

for *ρ_k_*(*x*) satisfying 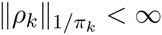. Thus, we have

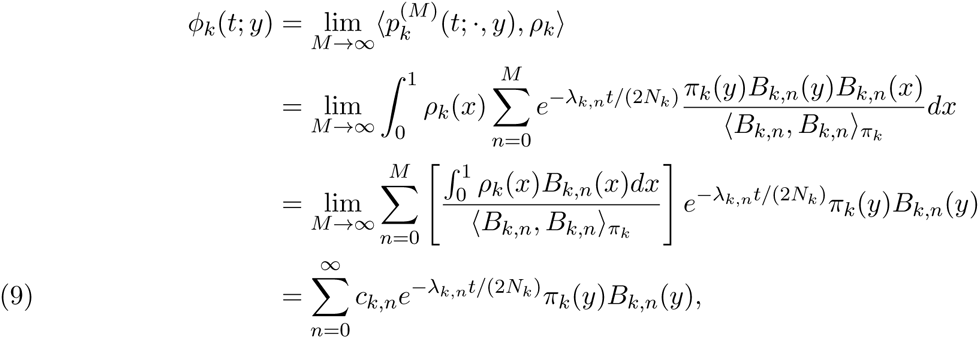

where

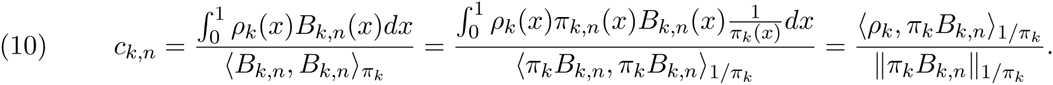

Determining the generalization, *ϕ_k_*(*t*; *y*), for any initial condition *ρ_k_* ∈ *L*^2^([0,1], 1/*π_k_*) thus amounts to projecting *ρ_k_* onto the functions 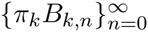 and plugging these coefficients into Equation (9) (Steinrücken *et al*. 2014, Vogl 2014b).

#### 2.2.1. *Computing the eigenfunctions* 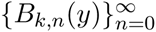.

Before discussing how to extend Equations (5) and (9) to the case of a population with piecewise constant parameters, which is the goal of this article, we first review the derivation of the eigenfunctions 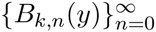 derived by Steinrücken *et al*. (2014).

Steinrücken *et al*. (2014) showed that the eigenfunctions 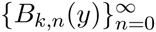 can be expressed as

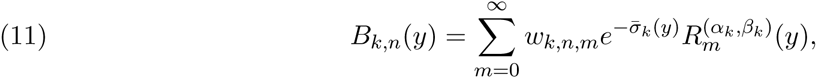

where 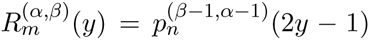, and 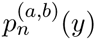 is the *n*th classical Jacobi polynomial. The vector *w_k,n_* = (*w_k,n_*_,0_, w*_k,n_*_,1_,…) is the left eigenvector of the matrix

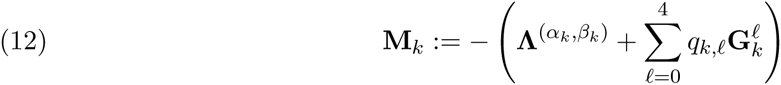

corresponding to the *n*th eigenvalue *λ_k,n_*. In Equation (12), 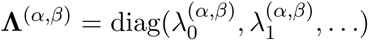 is the diagonal matrix with elements given by 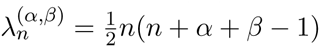 and *q_k_,_ℓ_* and 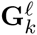 are given by

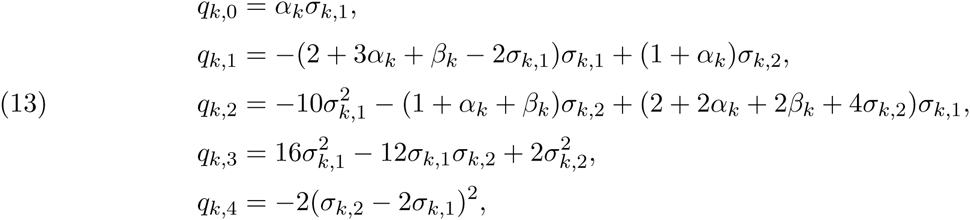

and

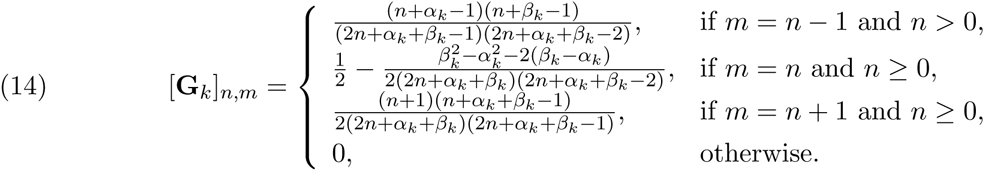

The eigenfunctions computed using Equation (11) can then be plugged into Equations (5) and (9), yielding the series expansions of *ϕ_k_*(*t*; *y*) and *p_k_*(*t*; *x*, *y*). In practice, we truncate the summations in Equations (5) and (9) at some large integer *N*, yielding

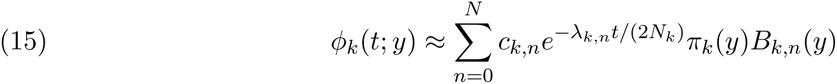

and

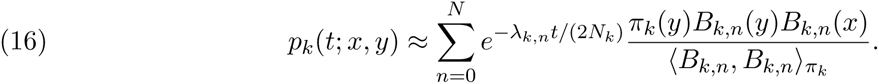

We also truncate the summation in Equation (11) at some large integer *M* ≥ *N*, yielding

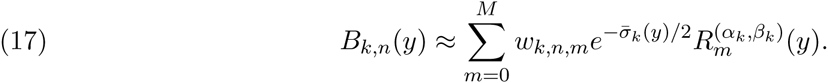

The eigenvalues, *λ_k,n_*, and eigenvectors, *w_k,n_*, of the matrix M*_k_* in Equation (12) are also computed by truncating the matrix M*_k_* to dimension *D* × *D*, for some large integer *D* ≥ *M*.

### 2.3. The TDF in a population with piecewise-constant parameters.

The goal of this work is to extend the results of Song and Steinrücken (2012) to populations with piecewise constant parameters. The challenge in computing the TDF for such piecewise constant populations lies in knitting together the expressions for the densities *ϕ_k_*(*t*; *y*) across the different epochs. This knitting procedure can be accomplished by taking the density at the end of epoch *k* as the initial condition for the density in epoch *k* + 1 (i.e., *ρ_k_*_+1_(*y*) = *ϕ_k_*(*t_k_* − *t_k_*_−1_; *y*)) by transforming the time-propagated coefficients 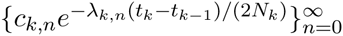 at the end of epoch *k* into the unpropagated coefficients 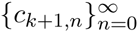 at the beginning of epoch *k* + 1.

Here, we focus on the extension of the generalization *ϕ_k_*(*t*; *y*) to multiple epochs, rather than generalizing *p_k_*(*t*; *x*, *y*) itself. The TDF, *p_k_*(*t*; *x*, *y*), along with generalized transition densities for other initial distributions *ρ_k_*(*x*) are obtained as special cases of *ϕ_k_*(*t*; *y*) in Section 3 by fitting the initial values of the coefficients 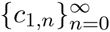 to different initial conditions, *ρ_k_*(*x*).

### 2.4. Transforming coefficients across epochs.

Plugging Equations (16) and (17) into the condition *ρ_k_*_+1_(*y*) = *ϕ_k_*(*t_k_* − *t_k_*_−1_; *y*) and dividing both sides by 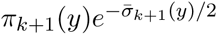 gives

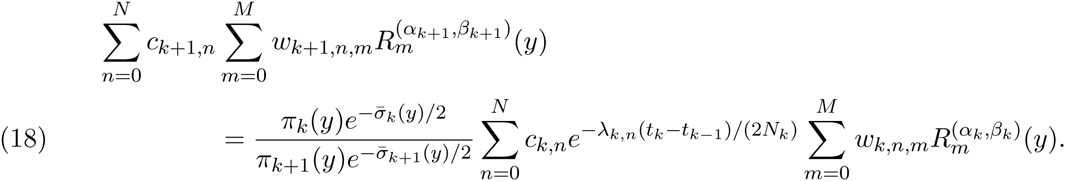

Because the left-hand side of Equation (18) is a polynomial of degree *M*, it is determined by *M* + 1 points. Therefore, we can determine the coefficients on the left-hand side by evaluating both sides of Equation (18) at a set of points y = {*y*_0_, …, *y_M_*}. We choose the set y to be the Chebyshev nodes because they minimize Runge’s phenomenon (Epperson 1987).

Evaluating Equation (18) at each of the points {*y*_0_, …,*y_M_*} and re-writing Equation (18) in matrix form gives

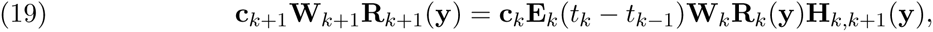

where the quantities in Equation (19) are given by

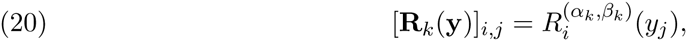

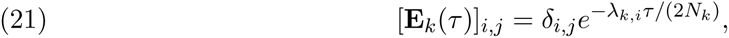

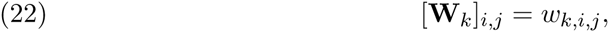

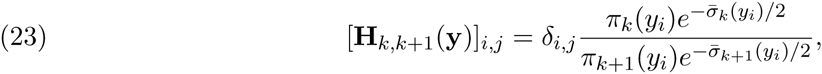

and **c***_k_* = (*c_k_*_,0_, *c_k_*_,1_,…). In Equations (21) and (23), *δ_i,j_* is the Kronecker delta function satisfying *δ_ij_* = 1 if *i* = *j* and *δ_ij_* = 0, otherwise. The coefficients **c***_k_*_+1_ in epoch *k* + 1 are obtained from the coefficients **c***_k_* in epoch *k* by solving Equation (19) for **c***_k_*_+1_ using standard approaches for solving linear systems.

The generalization, *ϕ_K_*(*t*; *y*), of the TDF at time 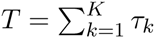 is evaluated by iteratively solving Equation (19) to obtain **c***_K_* in the final epoch, *K*, starting from a set of initial coefficients, **c**_1_. The final value of *ϕ_K_*(*t*; *y*) at time *T* is then computed as

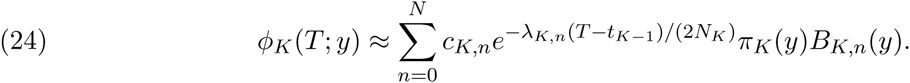

Equation (24) can be used to compute *ϕ_K_*(*t*; *y*), at any time *t* ∈ [0, *T*] by defining *T* = *t*, and choosing *K* to be the interval such that *t* ∈ (*t_K_*_–1_, *t_K_*].

## 3. Initial conditions

By fitting the coefficients, 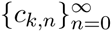, in Equation (24) to different initial distributions of frequencies at time *t* = 0, we can obtain the multi-epoch TDF, along with generalizations of the multi-epoch TDF to other initial distributions. Formulas for the starting coefficients **c**_1_ = (*c*_1,0_, *c*_1,1_,…) were presented in Steinrücken *et al*. (2014) for three different initial conditions: mutation drift balance, mutation selection balance, and the case of an initial starting frequency, *x*_0_. For completeness, these formulas are presented again below.

### 3.1. Initial frequency.

When the initial condition is a specified frequency, *x*_0_, the initial density is *ρ*_1_(*x*) = *δ*(*x* − *x*_0_), where *δ*(·) is the Dirac delta distribution. Steinrücken *et al*. (2014) showed that this choice of initial conditions gives rise to the transition density function *p*_1_(*t*; *x*_0_, *y*), where the coefficients 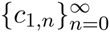 are given by

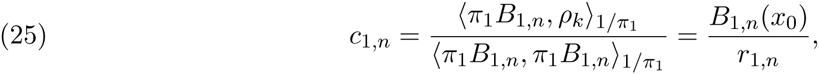

where

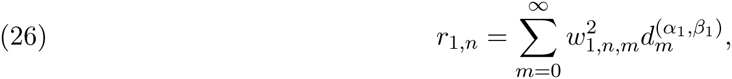

and

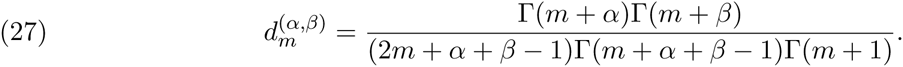

### 3.2. Mutation-selection balance.

Under the initial condition of mutation-selection balance, the initial density is the normalized stationary density

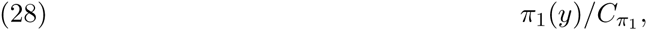

where *π*_1_(*y*) is given in Equation (6) and 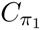 is a normalizing constant defined such that 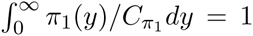. Using this initial distribution, Steinrücken *et al*. (2014) showed that the initial coefficients are given by

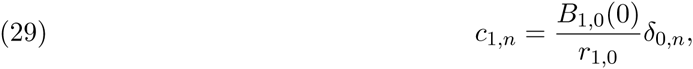

where *r*_1,0_ is given by Equation (26). The numerator in Equation (29) is given by

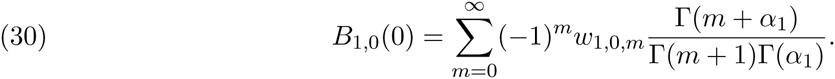

### 3.3. Mutation-drift balance.

Finally, under the initial condition of mutation-drift balance, the initial density is given by

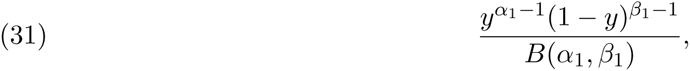

where *B*(*α*, *β*) is the beta function. Steinrücken *et al*. (2014) showed that the initial coefficients in the case of mutation-drift balance are given by

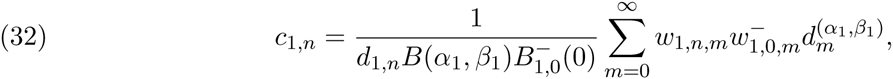

where the values 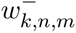 are the entries of the *n*th left eigenvector of the matrix M*_k_* with *σ*_1,1_ and *σ*_1,2_ replaced by − *σ*_1,1_ and −*σ*_1,2_, respectively, and 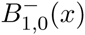 is the corresponding eigenfunction.

## 4. Implementation

Our algorithm has been implemented in JAVA. The inputs to the program are the effective population sizes (number of diploid individuals) ***N*** = (*N*_1_, …, *N_K_*); epoch durations ***τ*** = (*τ*_1_, …, *τ_K_*); per-generation mutation rates ***a*** = (*a*_1_, …, *a_K_*) and ***b*** = (*b*_1_, …, *b_K_*); selection parameters *s*_1_ = (*s*_11_, …, *s_K_*_1_) and *s*_2_ = (*s*_12_, …, *s_K_*_2_); initial allele frequency *X*_0_; and the time *t* ∈ [0, *T*] at which the TDF will be evaluated. A plot of the TDF evaluated at each epoch boundary point (*t* = *τ*_1_, *τ*_1_ + *τ*_2_, and *T*) in Figure 1 is shown in Figure 2. The full command options are detailed in the user manual distributed with the software.

**Figure 2.**
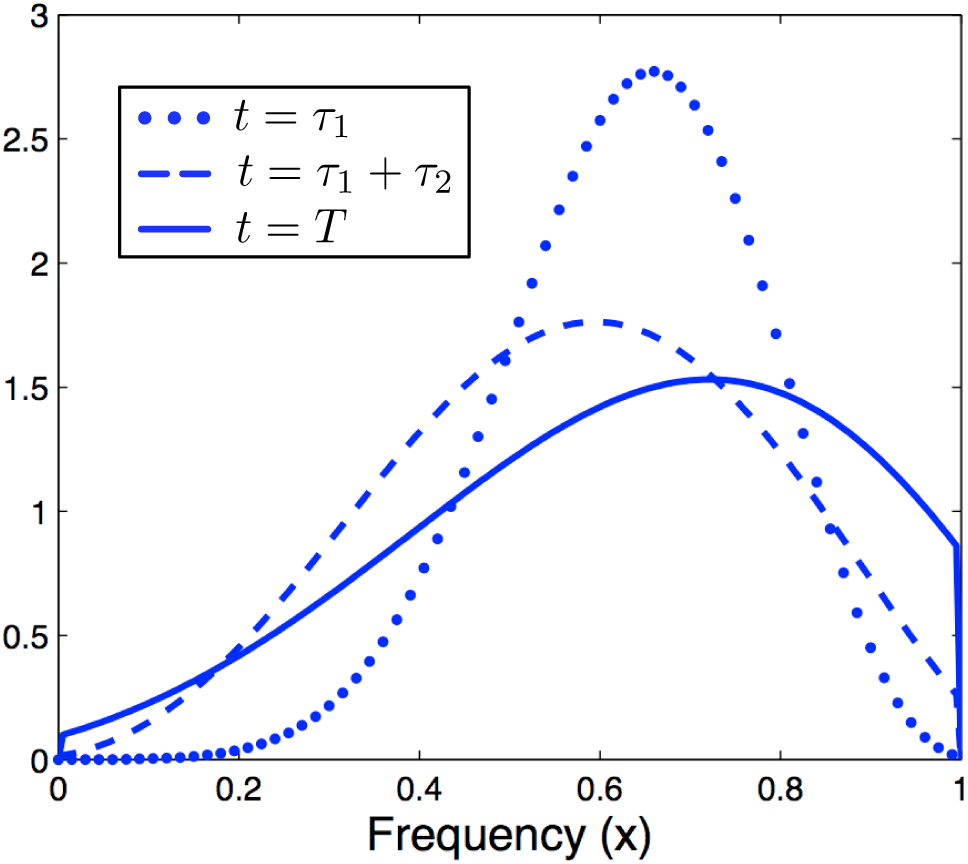
Plot of the TDF for the model shown in Figure 1 with the parameters specified in the example in Section 4, evaluated at the times *t*_1_, *t*_2_, and *T*.

The approach described in Section 2.3 for knitting together transition densities across epochs, combined with the method of Song and Steinrücken (2012) for computing the eigenfunctions (Section 2.2.1), produces a computationally efficient method for computing *p_K_*(*t*; *x*, *y*) and *ϕ_K_*(*t*; *y*). Table 1 shows the runtime of our method, SpectralTDF, for different numbers of epochs, and for different values of the parameters that control the precision of the eigenvalue computations.

**Table 1.**
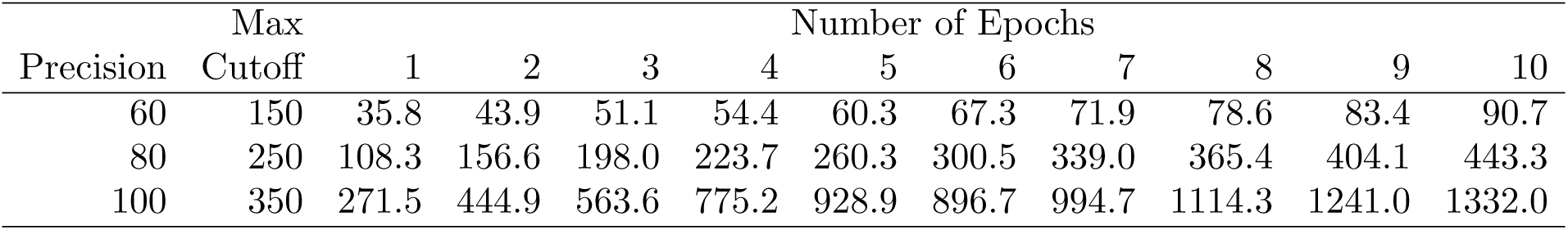
Runtime (in seconds) of the SpectralTDF algorithm for different numbers of epochs.

High precision computations are sometimes required when selection coefficients are large and waiting times between sampling events are short. However, such high precision computations are often unnecessary. Table 1 shows that SpectralTDF can be used to compute the TDF in a population with ten epochs in under two minutes, and for scenarios requiring higher precision in under ten minutes.

## 5. Discussion

Our implementation provides a fast and numerically stable method for computing the TDF for a general model with piecewise-constant population sizes and a broad range of time-varying mutation and selection parameters. It also allows for a variety of initial conditions, including a specified initial frequency and stationary distributions under mutation-selection balance or mutation-drift balance.

The JAVA implementation is designed to be used either as a stand-alone application or in combination with other methods. For example, the code can be easily incorporated into the method of Steinrücken *et al*. (2014), allowing the inference of selection parameters from time series data sampled from populations with time-varying demographic and selection parameters. In general, the method we present provides a flexible and efficient tool for studying the evolution of allele frequencies over time under complex evolutionary scenarios.

## Acknowledgement

We thank Daniel Živković and Anand Bhaskar for helpful discussions and collaboration on earlier related work. This research is supported in part by NIH grants R01-GM094402 (MS, YSS) and R01-GM109454 (EMJ), and by a Packard Fellowship for Science and Engineering (YSS).

## References

Barbour, A., Ethier, S., and Griffiths, R. (2000). A transition function expansion for a diffusion model with selection. Annals of Applied Probability, pages 123–162.

Bollback, J., York, T., and Nielsen, R. (2008). Estimation of 2*N_e_*s from temporal allele frequency data. Genetics, 179, 497–502.

Epperson, J. (1987). On the Runge example. American Mathematical Monthly, 94(4), 329–341.

Ewens, W. (2004). Mathematical Population Genetics: I, 2nd ed. Springer.

Griffiths, R. (1979). A transition density expansion for a multi-allele diffusion model. Advances in Applied Probability, pages 310–325.

Kimura, M. (1955). Stochastic processes and distribution of gene frequencies under natural selection. In Cold Spring Harbor Symposia on Quantitative Biology, volume 20, pages 33–53. Cold Spring Harbor Laboratory Press.

Kimura, M. (1957). Some problems of stochastic processes in genetics. The Annals of Mathematical Statistics, pages 882–901.

Shimakura, N. (1977). Equations différentielles provenant de la génétique des populations. Tohoku Mathematical Journal, Second Series, 29(2), 287–318.

Siepielski, A., DiBattista, J., and Carlson, S. (2009). Its about time: the temporal dynamics of phenotypic selection in the wild. Ecology Letters, 12(11), 1261–1276.

Song, Y. S., and Steinrücken, M. (2012). A simple method for finding explicit analytic transition densities of diffusion processes with general diploid selection. Genetics, 190(3), 1117–1129.

Steinrücken, M., Wang, Y., and Song, Y. S. (2013). An explicit transition density expansion for a multi-allelic Wright-Fisher diffusion with general diploid selection. Theor. Popul. Biol., 83, 1–14.

Steinrücken, M., Bhaskar, A., and Song, Y. S. (2014). A novel spectral method for inferring general diploid selection from time series genetic data. Annals of Applied Statistics, 8(4), 2203–2222.

Vogl, C. (2014a). Biallelic mutation-drift diffusion in the limit of small scaled mutation rates. arXiv, page 1409.2299.

Vogl, C. (2014b). Computation of the likelihood in biallelic diffusion models using orthogonal polynomials. Computation, 2, 199–220.

Živković, D., Steinrücken, M., Song, Y. S., and Stephan, W. (2015). Transition densities and sample frequency spectra of diffusion processes with selection and variable population size. Genetics, doi:10.1534/genetics.115.175265.

Waxman, D. (2011). A unified treatment of the probability of fixation when population size and the strength of selection change over time. Genetics, 188, 907–913.

Williamson, S., Hernandez, R., Fledel-Alonl, A., Zhu, L., Nielsen, R., and Bustamante, C. (2004). Simultaneous inference of selection and population growth from patterns of variation in the human genome. Proc. Nat. Acad. Sci., 102(22), 7882–7887.

